# Bacteriophages Playing Nice: Lysogenic bacteriophage replication stable in the human gut microbiota

**DOI:** 10.1101/2022.03.23.485530

**Authors:** Steven G. Sutcliffe, Alejandro Reyes, Corinne F. Maurice

## Abstract

1.1.

The human gut is a dense microbial community, of which bacteria and bacteriophages are the majority. Bacteriophages, viruses of bacteria, exist stably, without major fluctuations in the gut of healthy individuals. This stability appears to be due to an absence of ‘kill-the-winner’ dynamics, and the existence of ‘piggy-back-the-winner’ dynamics, where lysogenic replication rather than lytic replication occurs. Revisiting the deep-viral sequencing data of a healthy individual studied over 2.4 years, we were able to improve our understanding of how these dynamics occur in healthy individuals. We assembled prophages from bacterial metagenomic data and show that these prophages were continually switching from lysogenic to lytic replication. Prophages were the source of a stable extracellular phage population continually present in low abundance, in comparison to the lytic-phage population, where taxonomic diversity diverged over 2.4 years. The switch to lytic replication, or prophage induction, appears to occur mostly through spontaneous prophage induction. The observed phage dynamics of regular spontaneous induction are ecologically important as they allow prophages to maintain their ability to replicate, avoiding degradation and their loss from the gut microbiota.

**Significance Statement:** It has been eight years since Minot and colleagues published their landmark longitudinal study of phages in the gut. In the years following, the bioinformatic field improved in great strides, including the methods of bacterial-genome assembly, phage-identification, and prophage detection. We leveraged the unprecedented deep sequencing of phages in this dataset by adding bacterial assembly and prophage detection analyzes. We show clearly for the first time that ‘piggy-back-the-winner’ dynamics are maintained in the gut through spontaneous prophage induction, and not widespread triggered prophage induction. These dynamics play an important ecological role by creating a stable subpopulation of phages, which could help explain how phages are maintained over the 2.4 years timeframe that this individual was studied.

## 1.3. Introduction

The human gut is home to a diverse and abundant community of microorganisms that are central to human health and development. The most abundant members of this community are bacteria, found in the trillions, and bacteriophages (abbreviated phages), with abundances almost equal to bacteria [1]. Phage and bacterial communities co-evolve over the lifespan of the human host through a variety of infection dynamics [2] shaped by several factors: age, diet, medication consumption, and disease [2-4]. Years of host-bacteria-phage interactions during human growth and development result in an adult gut microbiota that is unique to each individual [5], with strongly correlated viral and bacterial communities [6]. The gut microbiota shows remarkable taxonomic stability, whereby bacterial diversity is a source of resilience to perturbations [7]. Understanding host-bacteria-phage interactions is of great importance to maintain, or improve, gut health.

Bacteria-phage relationships are driven by complex and dynamic interactions (see [8] for an overview). These interactions range from strictly parasitic to symbiotic, depending on the replication cycle undergone by phages (lytic and lysogenic, respectively) [2, 9]. The lysogenic replication cycle differs from lytic replication as the phage genome integrates into the bacterial genome as a prophage [2, 9]. Phages capable of lysogenic replication are referred to as temperate, and their bacterial host as lysogens. Both lysogens [10] and temperate phages [11, 12] have been observed in high abundance in the gut of healthy individuals. A longitudinal study of ten healthy adults also showed that the proportion of temperate phages varies between individuals, yet is relatively stable over time [5]. Understanding how lysogeny persists in the gut of healthy individuals will help contextualize the uniqueness of an individual’s gut virome [5], and the resulting viral-bacterial dynamics [6].

Lysogeny in the adult healthy gut has been hypothesized to stem from ‘kill-the-winner’ dynamics that play out during infancy [2, 13], leading to ‘co-adaptation as a means to stabilize the interactions between phages and hosts’ [14]. Co-adaptation has also been observed with CrAss-like-phages (e.g., ΦCrAss001) that co-exist with their *Bacteroides* host and persist in high abundance in the human gut [15] through a lytic-lysogeny intermediate [14, 16]. Dense populations of replicating bacteria [17, 18], as found in the gut, show increased rates of lysogeny through ‘piggyback-the-winner’ dynamics [19, 20]. Increased bacterial density increases the rate of lysogeny by phage coinfection [20], or through host-density regulated molecular switches [21]. Once integrated, prophages can provide a fitness advantage to their bacterial host through protection from further infection by superinfection exclusion or immunity [22], and the introduction of new genes encoding for virulence factors, antibiotic resistance, or novel metabolic functions [23]. In this case, prophages persist by ‘making-the-winner’ [20].

The benefits of lysogeny for bacteria are counter-balanced by the competitive cost of prophages being genetic elements that can switch to lytic replication through induction. Bacteria limit this switch by accumulating mutations within prophage regions at higher rates, rendering prophages inactive and incapable of lytic replication [24]. Hence, it is important to distinguish active prophages which can still be induced from inactive prophages. Prophage induction is typically triggered by extrinsic stimuli that result in DNA damage [25]. In the mammalian gut, external factors such as diet [26] and antibiotics [27] have been shown to trigger prophage induction. Prophages of gut bacterial isolates have been induced by dietary compounds [28], short-chain-fatty acids [29], antibiotics, and other medications [27]. Human pathologies, such as Crohn’s disease, could also lead to an increase in prophage induction [30]. Yet, prophage induction can also occur in the absence of external triggers, in a process referred to as spontaneous prophage induction [31]. In contrast with triggered prophage induction, spontaneous prophage induction leads to a small subset of prophages undergoing lytic replication [32] and is thought to be caused by intrinsic factors like stochastic gene expression or sporadic DNA damage [25].

We sought to determine how prophage induction contributes to the gut virome of healthy individuals. We hypothesize that most prophages in the gut are capable of active replication and replicate by regular spontaneous prophage induction. Spontaneous prophage induction is the most likely cause of prophage induction in the absence of disease, antibiotic use, or drastic dietary changes. Active prophages undergoing regular spontaneous prophage induction would translate to a small but stable fraction of extracellular phage population present in the gut.

To test our hypothesis and better understand the role of lysogenic bacteria and temperate phages in the gut, we revisited a previously published dataset of sequenced bacterial and viral metagenomic gut samples of a healthy individual over the course of 2.4 years [33]. This dataset was selected based on longitudinal sampling with sufficient resolution to observe prophage induction [34] and the detection of active prophages [26, 35]. We report that prophages contribute a stable, continuous source of free temperate phages in this healthy individual through spontaneous induction, while triggered prophage induction is rare and concerns only a few prophages. Our results suggest evolutionary or adaptive constraints between bacteria and phages in the gut that limit highly disruptive triggered prophage induction events in favour of spontaneous prophage induction.

## 1.4. Methods

### 1.4.1. Data Set

We used the previously published data of a healthy male, whose fecal samples were collected at sixteen time points spread over 884 days (∼2.4 years) [33]. The healthy twenty-three-year-old did not take antibiotics over the course of the experiment. The viral fraction was separated and sequenced at all sixteen timepoints, and eight timepoints were sequenced twice (Supplementary Figure 1A). Bacterial metagenomics were also obtained from fecal samples at three time points (once per week) during the same experimental time frame (Supplementary Figure 1A). For more details, see [33].

### 1.4.2. Viral Assembly

Sequence reads from viral-enriched libraries were trimmed with Trimmomatic V.0.36 [36], minimum quality 35 and minimum length of 70 (SLIDINGWINDOW:4:35 MINLEN:70 HEADCROP:10). As recommended [37], we assembled viral contigs for each sequence run separately with Spades [38] V.3.13.0 using the metaSpades option [39]. Viral assembled contigs were pooled, resulting in 291,758 contigs. Contigs less than 1kb in length were removed, resulting in 24,845 viral contigs. We used CD-HIT-EST V4.8.1 [40, 41] with 0.95 similarity threshold, 8-word size, 0.9 length cut-off to cluster the contigs from the different samples, resulting in 22,091 non-redundant viral contigs. We then selected for phage contigs, as those fulfilling at least one of the following three criteria: 1) Detected as viral by VirSorter (V.1.0.6) with custom database option additionally using the Gut Virome Database [42]; 2) at least three ORFs (predicted by Prodigal V.2.6.3 with metagenomic mode) with homology (HMMER V.3.1b2 hmmscan minimum e-value 1e-5) to PVOG database (Downloaded on Dec 1, 2020); or 3) BLASTn homology (e-value 1e-10, with 80% coverage of shortest contig) to Gut Virome Database [42]. This resulted in 14,444 phage contigs, of which 6,176 viral contigs were greater than 2.5kb in length.

### 1.4.3. Bacterial Assembled Genomes

Bacterial metagenomic reads were trimmed with Trimmomatic V.0.36 [36] (LEADING:3 TRAILING:3 SLIDINGWINDOW:4:15 MINLEN:36) and decontaminated for human sequences by aligning reads to *Homo sapiens* GRCh38 genome with bowtie2 [43] V.2.3.5.1. Remaining trimmed and decontaminated reads were pooled and assembled into contigs with megahit [44] V.1.2.7 using the default settings. We generated bacterial bins with contigs using MetaBat2 [45] V.2.12.1 -m 1500 (41 bins), CONCOCT [46] V.1.1.0 (77 bins) and MaxBin2 [47] V. 2.2.7 (14 bins). Bins were merged using DAS-Tool [48] V. 1.1.2. We used a score threshold of 0.35 to retrieve 27 bins. We then used CheckM [49] V.1.1.3 to confirm that all bins were unique. We selected bins that met the criteria of being either >40% complete and <10% contaminated by CheckM or having a DAS-Tool bin score of >0.4. This resulted in 25 medium-to-high quality bacterial bins. We assigned taxonomy to the bins using GTDB-Tk [50] V1.4.1 using the reference database [51] version r95. We determined the relative abundance as the percentage of reads that aligned to one of the 25 bins we detected. The total number of aligned reads per bin was normalized by bin size (Figure 1A).

**Figure 1.**
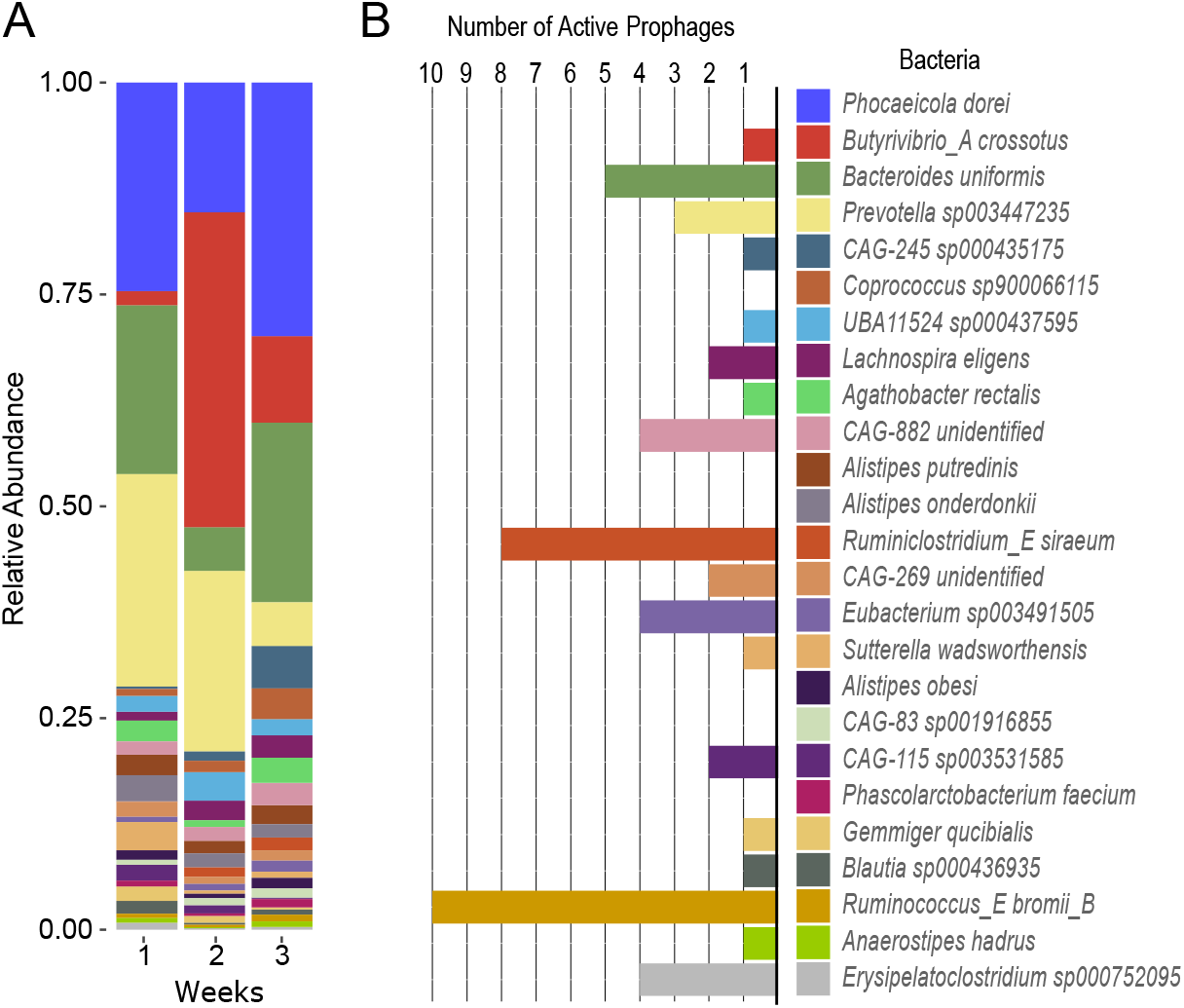
Distribution of bacterial lysogens in the gut of one healthy individual: (A) Relative abundance of all (25) medium-to-high quality assembled bacterial bins from metagenomic sequencing for each experimental week over the 2.4 years (B) Number of prophages determined to be active in each bacterial bin. The color-coding of bacterial bins is the same between the two panels.

### 1.4.4. Prophage Detection and Identifying Active Prophages

Prophages were detected within bacterial bins by combining various tools: PHASTER [52], VIRSorter (V.1.0.6) [53], VIBRANT (V.1.2.1) [54], PhageBoost (V.0.1.7) [55], and mVIR (V.1.0.0) [56]. We also used a custom alignment method, where we aligned the viral reads to each bacterial bin using bowtie2 (V.2.3.5.1), then used samtools mpileup to calculate coverage per base (with default perimeters). Using a sliding window of 1,000bp, if the average coverage was >10x, we considered the region as a possible prophage region. In total, we found 2,719 putative prophages. We merged prophage regions detected by multiple tools that overlapped by at least one base-pair resulting in 1,844 putative prophages. Only 17 prophages overlapped less than 100bp. Out of these 1,844 putative prophages, 651 prophages met one of our three phage criteria (see Viral assembled contigs methods). We then ran PropagAtE [57] (v1.0.0) on these remaining 651 detected prophages with a modified script (available upon request) that replaced host-coverage with the entire bin coverage when the flanking host region of the prophage was less than 5bp in length. It has been reported that prophages can be incorrectly binned when having multiple bins of closely related bacteria (in particular of the same species/genus) and when the prophage contig presents an absence of host-flanking regions [58]. We included prophages without host-flanking regions as the only bacterial bins assembled that shared genus (*Alistipes obesi, Alistipes putredinis*, and *Alistipes onderdonki*) did not contain any active prophages (Figure 1B). Finally, we used PropagAte’s criteria (default Cohen’s *d* test and prophage:host ratio) to identify 52 predicted active prophages in this dataset [57].

### 1.4.5. Viral Community

The viral population used in this study consisted of 6,176 assembled phage contigs and 52 active prophages. Out of the 6,176 assembled phage contigs, we removed 338 that had homology to the set of active prophages (BLASTn e-value 1e-5), resulting in a total of 5,890 non-redundant viral contigs. Quality trimmed reads were aligned to viral contigs using bowtie (V.2.3.5.1). Read coverage was normalized by sample using DESEQ2 V1.30.1 [59], then by viral contig length. Viral contigs were considered ‘present’ in a sample if their genome was covered in 75% of the length by at least 1x coverage [37]. Family-level taxonomic classification was performed by using a voting-approach after comparing genes on the amino-acid level against the viral subset of TrEMBL by Demovir (github.com/feargalr/Demovir). No CrAss-like-phages were predicted, and their absence was confirmed through additional comparisons of our viral contigs against Guerin crAss-like phage genomes [15], through BLASTn similarity and shared viral clusters (using VCONTACT2 [60]). Less than 0.05% of viral reads aligned to the Guerin crAss-like phage genomes [15], indicating a low-abundance of CrAss-like-phages in this individual.

Before predicting the replication strategy of viral contigs we selected the high-quality viral contigs (i.e., >= 40% complete and classified as high-quality by CheckV [61](V.0.7.0.)). These high-quality viral contigs (557) represent a mean of 82% (std. dev. 7.94) of quality controlled viral reads. We used Bacphlip [62] V.0.9.6 on our high-quality viral contigs to predict which were temperate (that is with a >50% chance of being temperate). Bacphlip is designed to be used on complete genomes and will under-report temperate phages when applied to incomplete or fragmented phage genomes.

## 1.5. Results

To study the prophages present within the genomes of bacterial lysogens, we used whole community metagenomic sequences and assembled 25 medium-to-high quality bacterial bins. All bacterial bins were taxonomically identified at the genus level, and 23 at the species level. The assembled bacterial community consisted mostly of Firmicutes and Bacteroidetes and one Proteobacteria, Sutterella wadsworthensis, a commonly found gut bacteria. These bins represent approximately 46%, 56%, 54% of metagenomic aligned reads on days 182, 852, 881, respectively. Bacteria not represented in the bins were likely at too low abundance for assembly and binning of adequate quality bins for prophage detection. Four of the bacteria represented between 67-79% of the mapped reads: Prevotella sp003447235, Phocaeicola dorei, Bacteroides uniformis, and Butyrivibrio_A crossotus (Figure 1A). Our bacterial diversity and bacterial bin abundances data are in line with what was reported in the original work using read-based methods [33].

Each bacterial bin was investigated for prophages using multiple tools (see Methods). Most prophages detected were detected by several tools (Supplementary Figure 2A), which led to the detection of 651 non-redundant putative prophages (Supplementary Figure 2B). We used Propagate [57] to separate prophage-like artifacts, or prophages no longer capable of replicating, from true prophages on the assembled bacterial bins. The prophages that were found to be actively replicating in our samples were deemed as ‘active prophages’. Of these 651 putative prophages, we found 52 active prophages (Supplementary Figure 2C). We excluded non-active prophages from this study as differentiating non-active prophages that are still capable of replication (true-positives) from prophage-like artifacts (false-positives) is not possible with our liberal prophage detection approach. We quantified that most bacteria (72%) contain at least one active prophage, and 40% of these bacterial lysogens contain multiple active prophages (Figure 1B).

We aligned all the viral metagenomic reads to see when each active prophage was found over the 2.4 years (Figure 2). Thirty eight (73%) of the active prophages were detected in the viral metagenomes (presence determined as in [37]) (Figure 2), further confirming their status as ‘active prophages’ (56% of bacteria contain at least one active prophage found in the viral sequenced fraction). More than half of active prophages were found at 9 separate time points and three different weeks (Figure 2). Over the 2.4 years, 19 prophages significantly increased in abundance during at least one time-point compared to the other time points (DESEQ2, adjusted p-value <0.05). The significant increase in abundance indicates a possible triggered prophage induction event. Eight prophages within six bacterial lysogens reached DESEQ2 normalized coverage 100x (Figure 3A), and of those, five were significantly increased (z-score >1.96 of log-transformed coverage) at one time point. Increased coverage occurred almost entirely during week 3, between days 881-885 (Figure 3A). *CAG-115 sp003531585* prophage 1 was the exception, as it rose to significant abundance during week 1 (day 184) as well as week 3 (day 881) (Figure 3A). Significantly increased prophages at week 3 were found in five different host-bacteria belonging to both Firmicutes and Bacteroidetes; meaning the stimuli triggering prophage induction are not phyla-specific and likely external ones. This contrasts with week 1 where triggered prophage induction was a species-specific event. Active prophages were found in low abundance, with a cumulative abundance typically less than 0.5% of the community, with the exception of day 881, which rose above 1% (Figure 3B). The continuous low-abundance of active prophages (Figures 3A-B), and continuous presence of active prophages over the 16 sampling times (Figure 2) together support a model of regular spontaneous induction. Triggered prophage induction on the other hand is limited to fewer time points (4 of 16 timepoints) and small fraction of lysogens (5 of 18).

**Figure 2.**
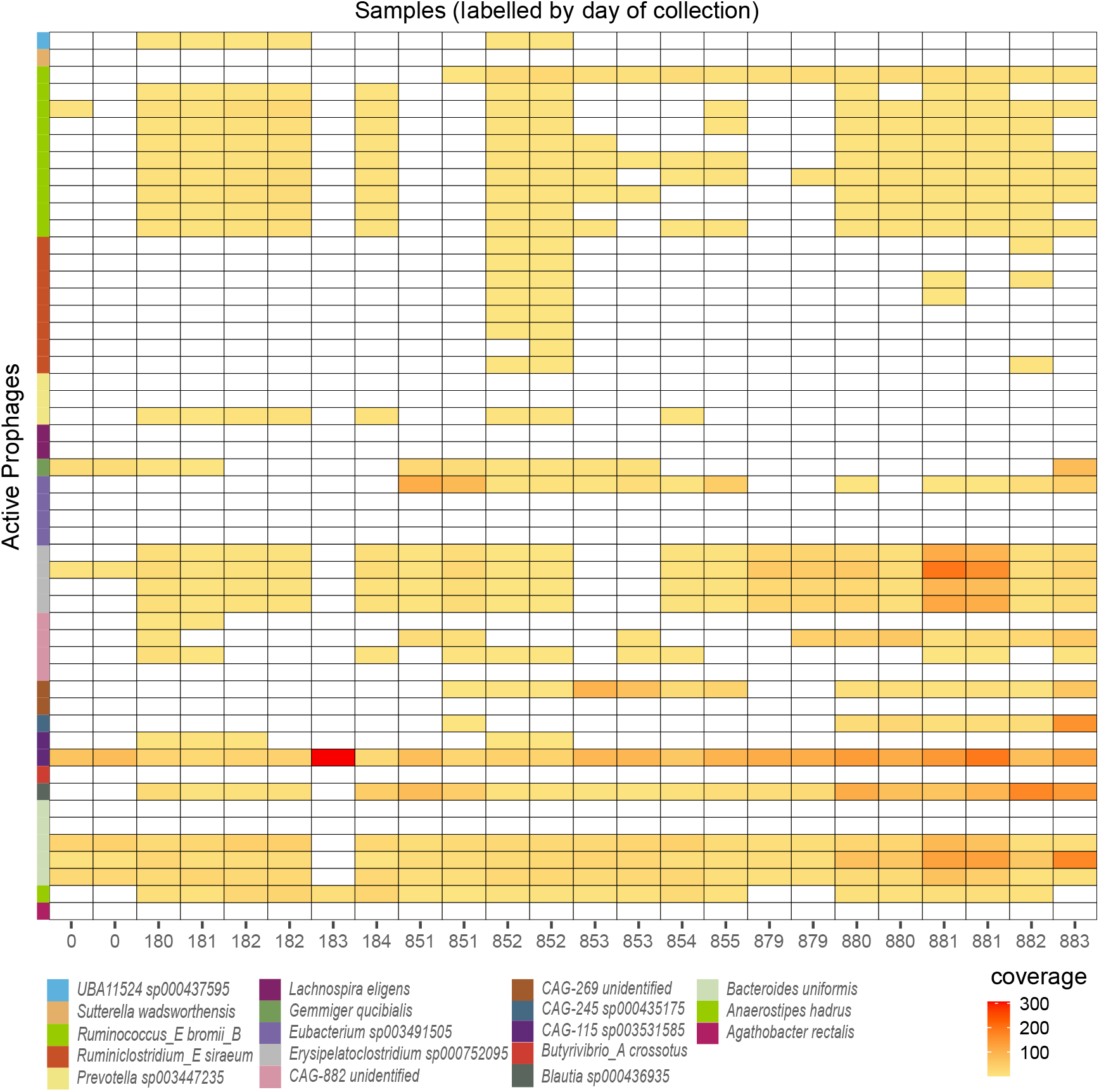
Presence of active prophages in the gut of one individual over 2.4 years: Heat map showing the relative abundance (coverage) of active phages (rows) in each of the viral-enriched sampled time points. White time points indicate an absence of prophages detected in sample. Presence was defined as viral reads covering prophage regions by at least 1x for more than 75% of prophage length in the sample. A total of 38 out of 52 active prophages were considered present in at least one time point. Normalized coverage of active prophages is displayed on timepoints when an active prophage was present in the sample.

**Figure 3.**
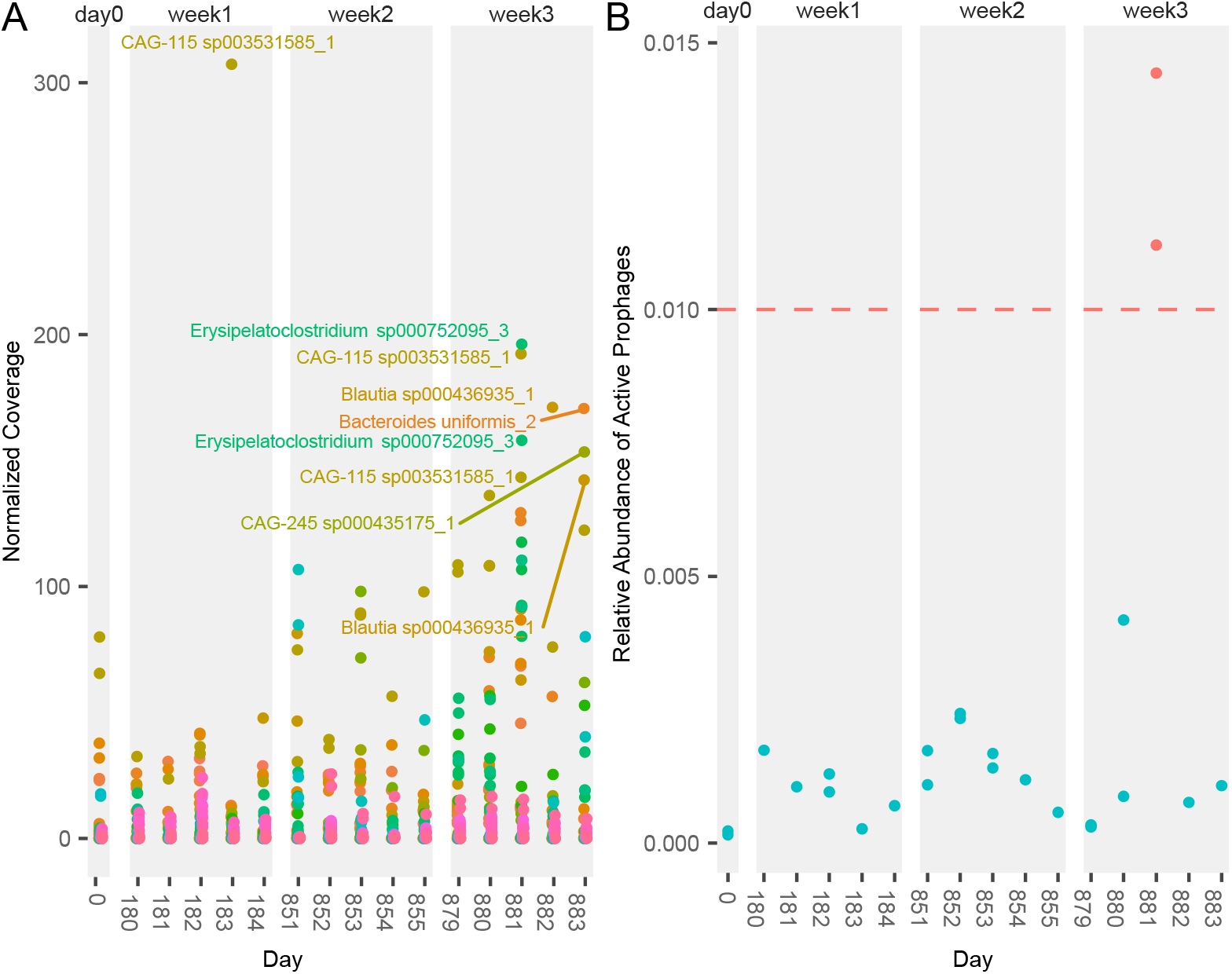
Spontaneous induction of prophages in a healthy individual dominates in comparison to rare, triggered prophage induction events: (A) Normalized coverage of each active prophage over the course of the experiment. Active prophages with significantly increased coverage (z-score > 1.96) are labeled. (B) Relative abundance of all active prophages in the virome fraction. Red time points are above the 1% relative abundance (dashed line).

From the viral metagenomic reads we assembled 6,176 phage contigs. We combined the 52 prophages to our analysis and removed 338 viral metagenomic assembled phage contigs that had homology to the set of active prophages, resulting in a total of 5,890 non-redundant viral contigs. We were able to classify 44% of these phage contigs taxonomically at the family-level (Supplementary Figure 3). A large percentage of phages at weeks 2 and 3 were unknown (Supplementary Figure 3). The relative abundance of classified phages at the family-level shows most members belonging to the *Microviridae* family (Figure 4A), which contrasted with the absence of CrAss-like-phages. CrAss-like-phages have also been observed to be at low-abundance in individuals with a high-abundance of *Microviridae* phages [5]. At the family-level, the viral community appears to be stable over the 2.4 years. However, we see an expansion of families belonging to *Caudovirales* (*Siphoviridae* and *Myoviridae*) at day 881 (Figure 4A) which corresponds to the increase in active prophages (that belong mostly to the *Siphoviridae* and *Myoviridae* families) from triggered prophage induction (Figure 3). In contrast, triggered prophage induction at day 183, which only impacted *CAG-115 sp003531585*, did not have an impact on the viral diversity. Triggered prophage induction, unlike spontaneous prophage induction, can significantly and rapidly alter the phage community diversity. Here, the effect of triggered prophage induction is transient, only lasting one day as *Microviridae* phages return to their high relative abundance the following day (day 882).

**Figure 4.**
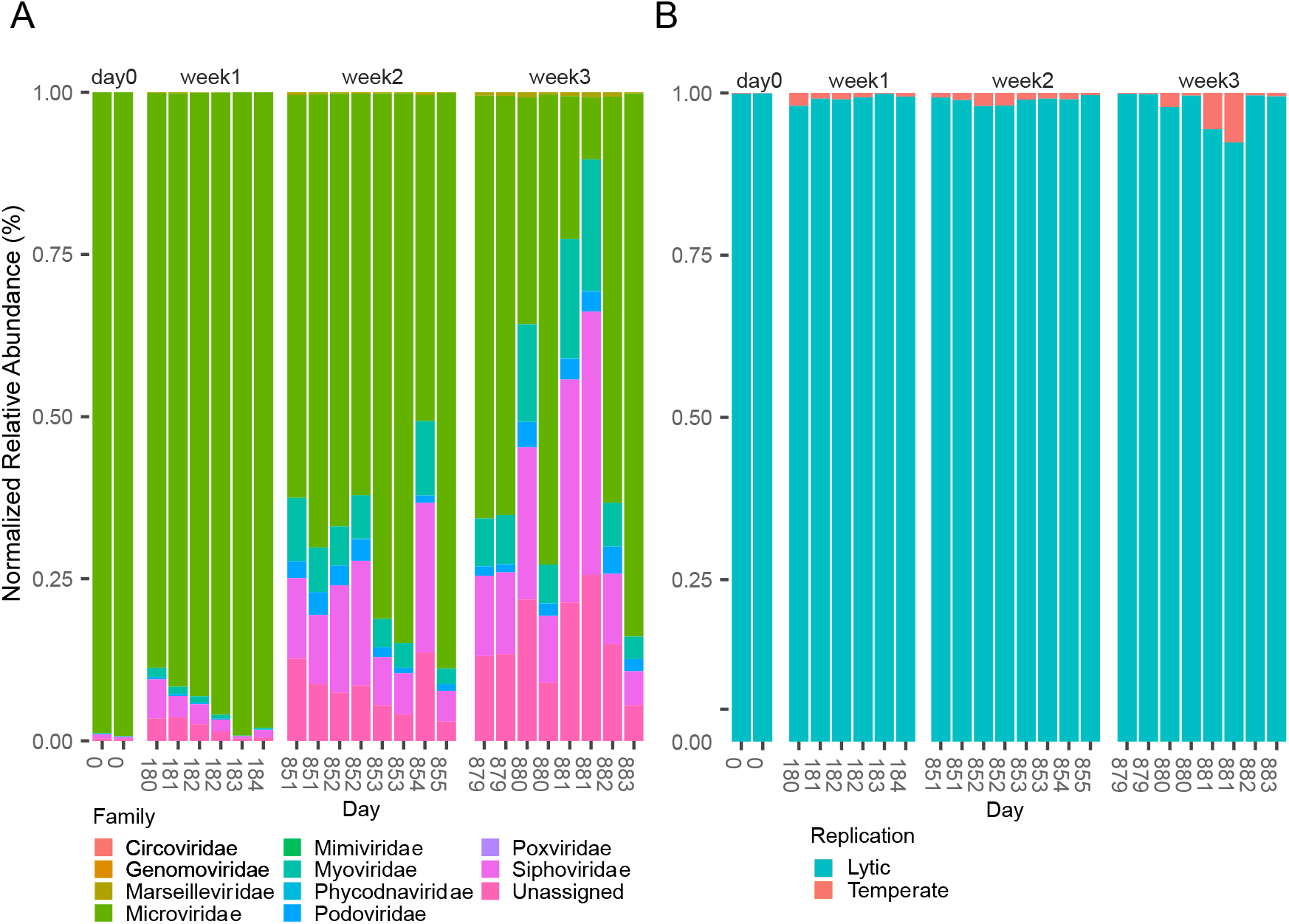
Diversity of phages in the gut of a healthy individual over 2.4 years: (A) Relative abundance at the phage family-level showing disruption of stability at day 881 by triggered prophage induction (B) Relative abundance of subset of high-quality viral contigs predicted to be either temperate or strictly lytic.

To determine if active prophages influence the proportion of temperate phages in the gut, we determined which of our phage contigs are potentially temperate using Bacphlip [62]. Lysogenic replication cycle prediction by Bacphlip is designed to be run on complete phage genomes, as incomplete genomes underestimate temperate phages due to an absence of genetic hallmarks. For temperate phage analysis, we took a subset of our phage contigs that were high-quality (>90% complete) or complete by CheckV [61]. This resulted in 557 phage contigs that represented a mean 82% (std. dev. 7.94) of viral reads. We found few temperate phages in high relative abundance (Figure 4B). However, at week 3, there is an increase of the temperate fraction from an average 1.3% to 2.1%, and peaking at day 881 (6.5%) (Figure 4B). This once again supports the idea that triggered prophage induction occurred at week 3, temporally altering the phage community.

To better understand how frequent spontaneous prophage induction and rare triggered prophage induction events shape the stability of phage community over time, we looked to discriminate between different patterns of community composition change (stochastic variation, directional change, and cyclical dynamics) using the approach of Collins *et al*. [63]. We investigated the change in phage taxonomic composition change with beta-diversity (Bray-Curtis dissimilarity) between time points of the whole phage community (5,890 phage contigs) over time. The regression line was significant (*p* = 2.2e-16) with a positive slope (0.025, adj. R-squared 0.9) indicating the community is diverging over time. In comparison, the active prophages are stable and significantly less divergent (*p* = 1.36e-05) with almost no positive slope (0.003, adj. R-squared 0.038), indicating the active prophages are more stable than the whole phage community (Figure 5AB). To confirm that the results from active prophages are not a sub-sampling artifact, we randomly sub-sampled 52 viral contigs from the whole community (20 iterations), and all were more divergent than active prophages. The rate of divergence over 2.4 years leads to three compositionally distinct viral communities that significantly grouped together by week (PERMANOVA Pr(>F) 0.001: day 0/week 1 (days 0-184), week 2 (days 851-855), and week 3 (days 879-883), with later groups clustering closer due to temporal proximity (Figure 5C). The stable active prophage population clusters less by week (PERMANOVA Pr(>F) 0.11) (Figure 5D). The slower divergence rate in active prophages leads to less separation by week, despite a higher baseline dissimilarity (0.58 for active prophages compared to 0.18 for the whole community). Active prophages thus appear to be a stable community of phages where composition is maintained over long periods of time, despite prophage induction events.

**Figure 5.**
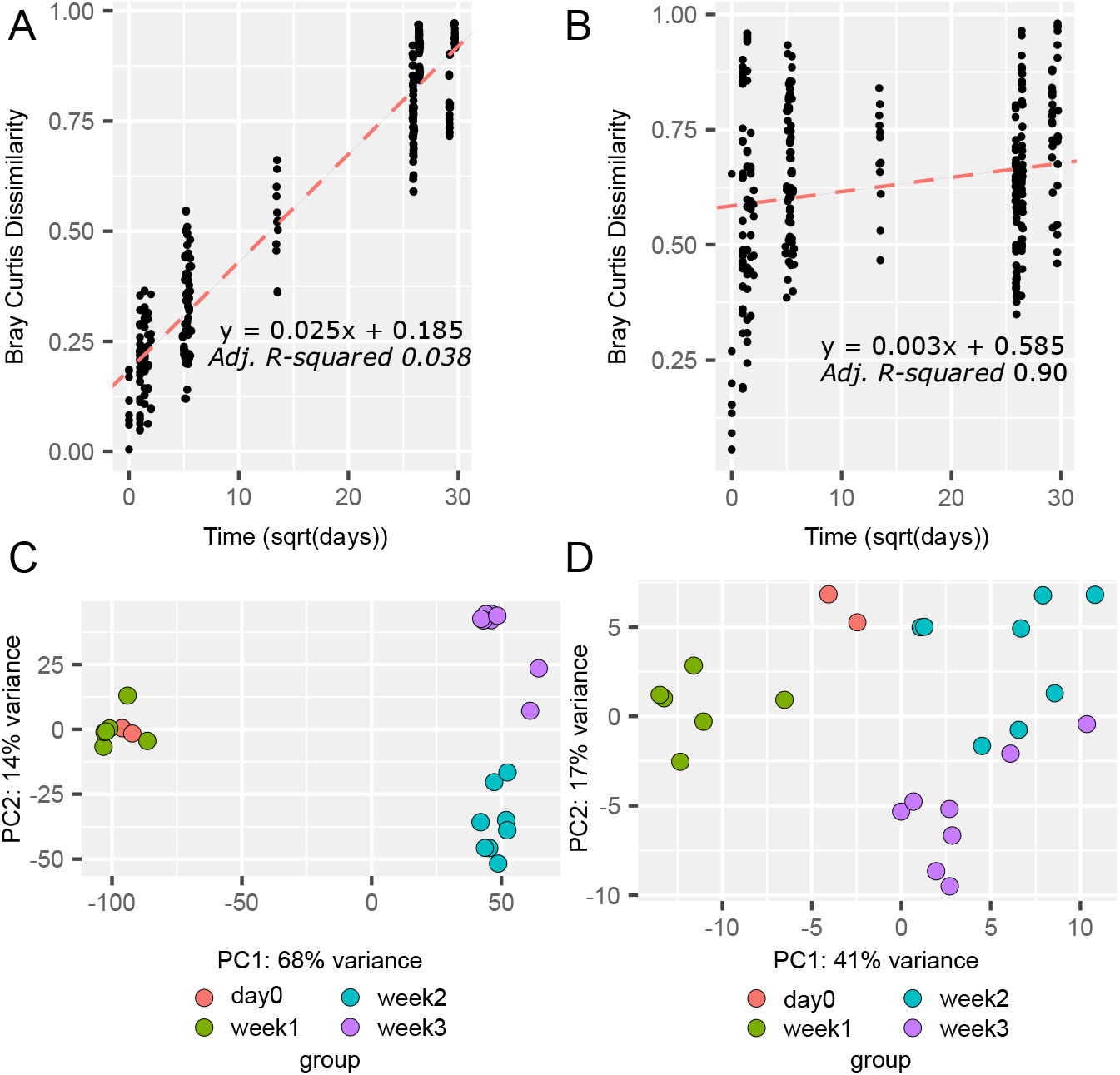
Stability of both viral and prophage communities over 2.4 years: Bray-Curtis Dissimilarity between samples by time elapsed distance to track the degree the rate of community change (slope of dotted line) in both the (A) whole viral community (p = 2.2e-16) with positive slope (0.0245), and linear (adj. R-squared 0.9) and the (B) prophage community (p = 1.36e-05) with a positive slope closer to zero (0.003) (C). PCA plot of DESEQ2 relative abundance of samples by time in the whole viral community (D) and in the prophage community.

We confirm that active prophages are commonly found in the genomes of gut bacteria, and that they replicate continually through spontaneous prophage induction. In the 16 time points sampled over 2.4 years, only four time points showed signs of triggered prophage induction. Triggered prophage induction at day 881, which involved multiple lysogens, was able to disrupt viral composition at a family-level and increase the abundance of temperate phages.

## 1.6. Discussion

By studying both the bacterial and viral communities of a healthy individual over the course of 2.4 years, we were able to link prophages to the extracellular phage community. We found that active prophages represent a taxonomically stable fraction of the phage community, present throughout the 2.4 years at low abundance. Triggered prophage induction involved only a small fraction of lysogens with active prophages (5 of 18). Only one lysogen underwent triggered prophage induction during week 1 (Figure 3A). This event did not have an impact on the phage community (Figure 4A), as there was no shift in phage taxonomic composition or change in the relative abundance of prophages (Figure 3B) or temperate phages (Figure 4B). In contrast, during week 3, more lysogens underwent triggered prophage induction, leading to a disruption of the phage community (Figure 4A), and relative abundance of prophages (Figure 3B), and temperate phages (Figure 4B). For the other 12 time points, prophage abundances support events of spontaneous prophage induction. The ecological consequence of this dynamic was a stable prophage fraction of the community (Figure 5BD) when compared to the rest of the community (Figure 5AC). Spontaneous prophage induction could be a source of stability in the phage gut community or result from the absence of perturbations.

Prophages are commonly found in the genomes of bacterial isolates [64], as well as bacteria found in the guts of humans [65] and mice [26]. We show that the gut microbiota of this healthy individual contains numerous active prophages (Figure 1B), similar to what was observed in the gut of healthy mice [26]. We leveraged multiple prophage predictors [66] and focused on medium-to-high quality bacterial bins, as well as prophages found on bacterial scaffolds without host-flanking regions [26]. Prophages from scaffolds without host-flanking regions decrease accuracy of prophage assignment to host [58], but was a necessary approach as few prophages were assembled with host flanking regions. In addition, few assembled bacteria belonged to the same genus, thus reducing the likelihood of mis-assigning hosts. When the prophages in the mouse gut [26] were revisited using only prophages with flanking host regions, there were less active prophages observed [67] than in the original study [26]. We confirmed that most bacteria contained active prophages, through their detection in the viral-sequenced fraction. Detection of prophages in the viral sequenced fraction despite their low abundance, is possibly due to the high sequencing depth of the original study. This allowed for the detection of spontaneously induced prophages. Future studies might benefit from hybrid assemblies of short-read and long-read sequences to improve the quality of bacterial metagenomic assembled genomes, and prophage prediction [68] so that all prophages can be assembled with host-flanking regions.

It has been argued that prophages, especially active prophages, represent a fitness cost to bacteria, as prophages represent extra genetic cargo that can act as a ‘molecular time-bomb’ [69], and therefore should be under selective pressure to be rendered inactive [24]. Despite this, we see most bacteria present here have active prophages in their genomes (Figures 1B, 2). This individual was sequenced well-above saturation [33], which allowed us to track low-abundance phages, and suggest that active prophages are undergoing spontaneous prophage induction and not triggered prophage induction (Figure 3 AB, 4AB). The differentiation between the two types of prophage induction offers an explanation as to how temperate phages might mitigate the fitness cost of actively replicating (reviewed in [25]). Spontaneous prophage induction would result in less bacterial death compared to triggered prophage induction (Supplementary Figure 1C), and prophage release could act as ‘bacterial warfare’ to closely related bacteria vulnerable to infection [70]. This process ultimately promotes lysogeny in the long-term [70]. These results support the hypothesis that spontaneous prophage induction is a mechanism by which prophages maintain their ability to remain active over long periods of time [25, 71].

The gut virome of a healthy individual is considered stable over time. At the family level, this individual’s gut virome was stable (Figure 4A), but at the contig level it was undergoing directional change (Figure 5A). The family-level stability was altered temporarily by a triggered prophage induction event at day 881, but not at week 1 (Figure 4A). Phage contigs in this data set, which represent more closely phage species or strains, were previously reported to be stable, as 80% of phage contigs were found at day 0 and day 883 (the end of the study) [33]. These findings relied on the Jaccard Index, focusing on the number of shared contigs, whereas we defined stability using Bray-Curtis dissimilarity, including contig relative abundance as well as presence/absence between time points. In doing so, we found that phage community composition is diverging (Figure 5A) [63]. Original findings focused on the most abundant phage contigs through manual curation, and since, progress has been made to automate the identification of phage contigs using command-line tools (e.g., VirSorter [53]), annotated phage protein, (e.g., pVOG [72]), and databases of gut viruses [42]. These improvements allowed us to include rare phage contigs, which increase dissimilarity. Active prophages undergo less compositional divergence over time than expected from the whole phage community, even with prophage induction. The decreased divergence is likely in part due to slower mutation rates of temperate phages compared to lytic phages [33], and not just spontaneous prophage induction.

We did not identify any CrAss-like phages in this individual, probably because CrAss-like phages have been shown to be in high abundance when temperate phages and *Microviridae* phages are in low abundance [5], which is not the case here. Lysogenic replication fits into a larger category of passive replication which includes pseudolysogeny and chronic infection [9]; an important characteristic for success in the gut, as this is how CrAss-like phages appear to be replicating [15]. Interestingly, it is a feature specific to adults, as the infant gut undergoes rapid changes where prophage induction appears to play a more important role [13, 73]. Passive-replication in the adult healthy gut might be the consequence of continuous ‘kill-the-winner’ dynamics that occur during infancy [2, 13]: over time, this pressure drives phages and bacteria to co-adapt, leading to increased stability between bacteria and phages in healthy adult gut [5, 14].

Numerous factors can alter both bacterial and viral compositions in the gut, including age, disease, drugs, diet, *etc*. [3, 4]. These factors are also capable of triggering prophage induction. It appears that in the absence of disease or antibiotics, this individual had a triggered prophage induction event at week 1 and 3. At week 3, prophage induction impacted multiple bacteria belonging to both Bacteroidetes and Firmicutes phyla, it is thus likely that some environmental factor is responsible. In contrast, the triggered prophage induction detected at week 1 seems more targeted and could be a species-specific event. Unfortunately, no metadata was collected for this study, so we hypothesize that non-antibiotic medication consumption [27] or a switch in diet [26] are likely responsible. We will need to test these hypotheses moving forward to have a better understanding of prophage induction in the gut. Currently, we do not know how species-specific triggers differ from community-level triggers of prophage induction. To test these hypotheses, large-scale studies with comprehensive metadata are needed [74] or implementing gnotobiotic mouse models to explore features causing prophage induction of gut prophages *in vivo* [35].

Longitudinal studies are important when studying the gut microbiota [75], including the virome. With daily sampling of this individual, we could detect triggered prophage induction occurring at day 881, but changes in the phage community were undetectable at days 880 or 882, or the rest of the week (Figure 3B, 4AB). This highlights how detecting prophage induction at the community level is difficult [76], and therefore using time-series of bacterial and viral metagenomics can provide insight into how active prophages contribute to extracellular intestinal phage communities [26]. Daily sampling of the gut community is necessary to detect triggered prophage induction [77] with sequencing depth above saturation [33]. This scale of daily observations is important to increase our understanding of phage-bacteria dynamics in the gut despite the challenges of increased sampling and cost.

## 1.7. Conclusion

Our work supports the hypothesis that lysogeny is a stabilizing force between bacterial and phage communities in the gut [14]. Prophages in the gut of this healthy individual appear to be balancing the benefits of stable integration with the risk of inactivation through regular spontaneous prophage induction. As phages undergo divergent evolution in the gut, we speculate that lysogeny offers a refuge from genetic divergence. Bacteria balance the benefits of accumulating prophages against the costs of having extra genetic cargo that can trigger cell lysis. Regular triggered prophage induction would increase the fitness cost of harbouring prophages and increase selective pressure for prophage inactivation. In conclusion, bacteria and temperate phage balance competing priorities to form a stable equilibrium in the gut and play nice.

## Notes

### Competing Interest Statement

The authors have declared no competing interest.

