## Supplementary Figures for "Bacteriophages Playing Nice: Lysogenic bacteriophage replication stable in the human gut microbiota"

### Supplementary Material

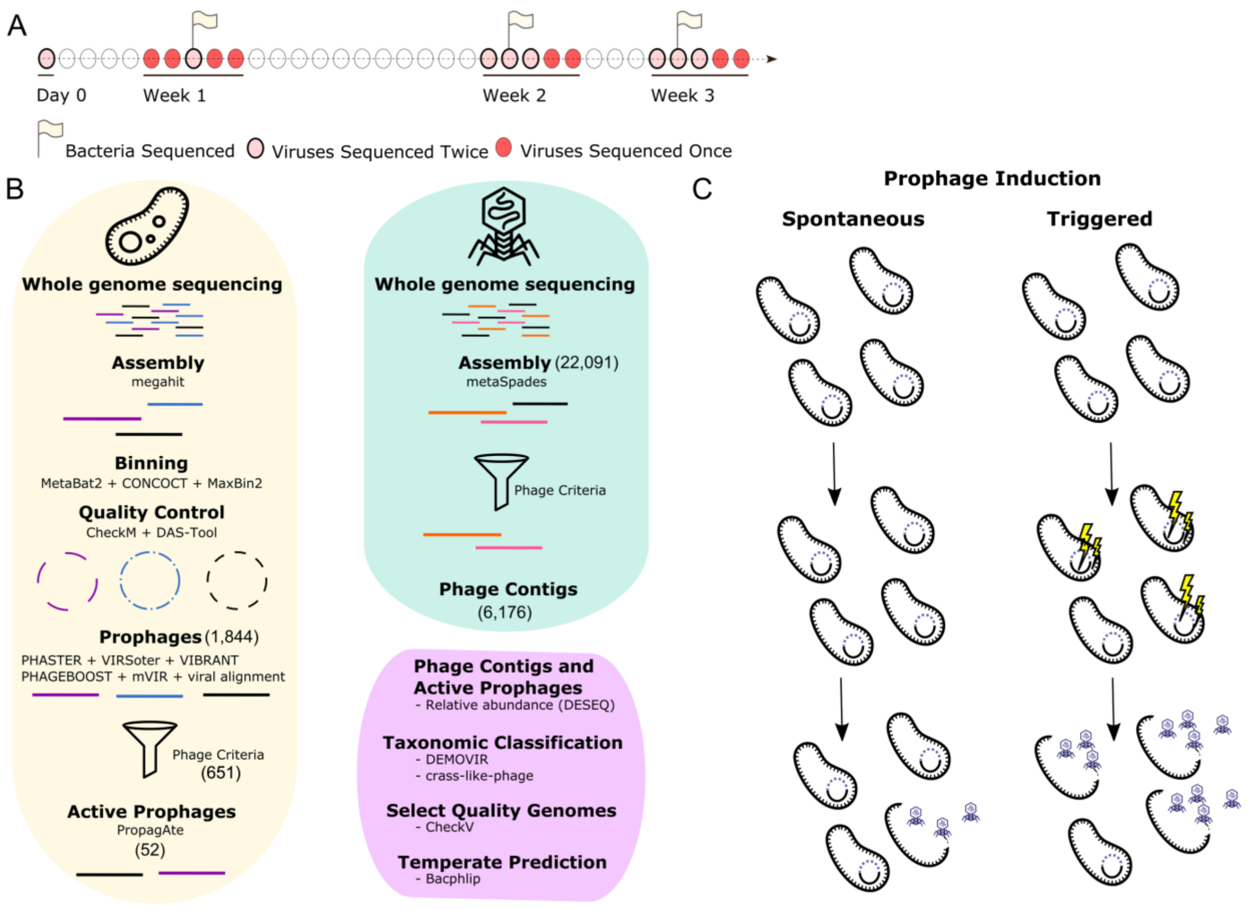


#### Supplementary Figure 1 Study Summary:

(A) Explanation of the sampling for the study (B) Methodology (C) Comparison of spontaneous prophage induction to triggered prophage induction. Lightning bolts represent external prophage induction triggers.

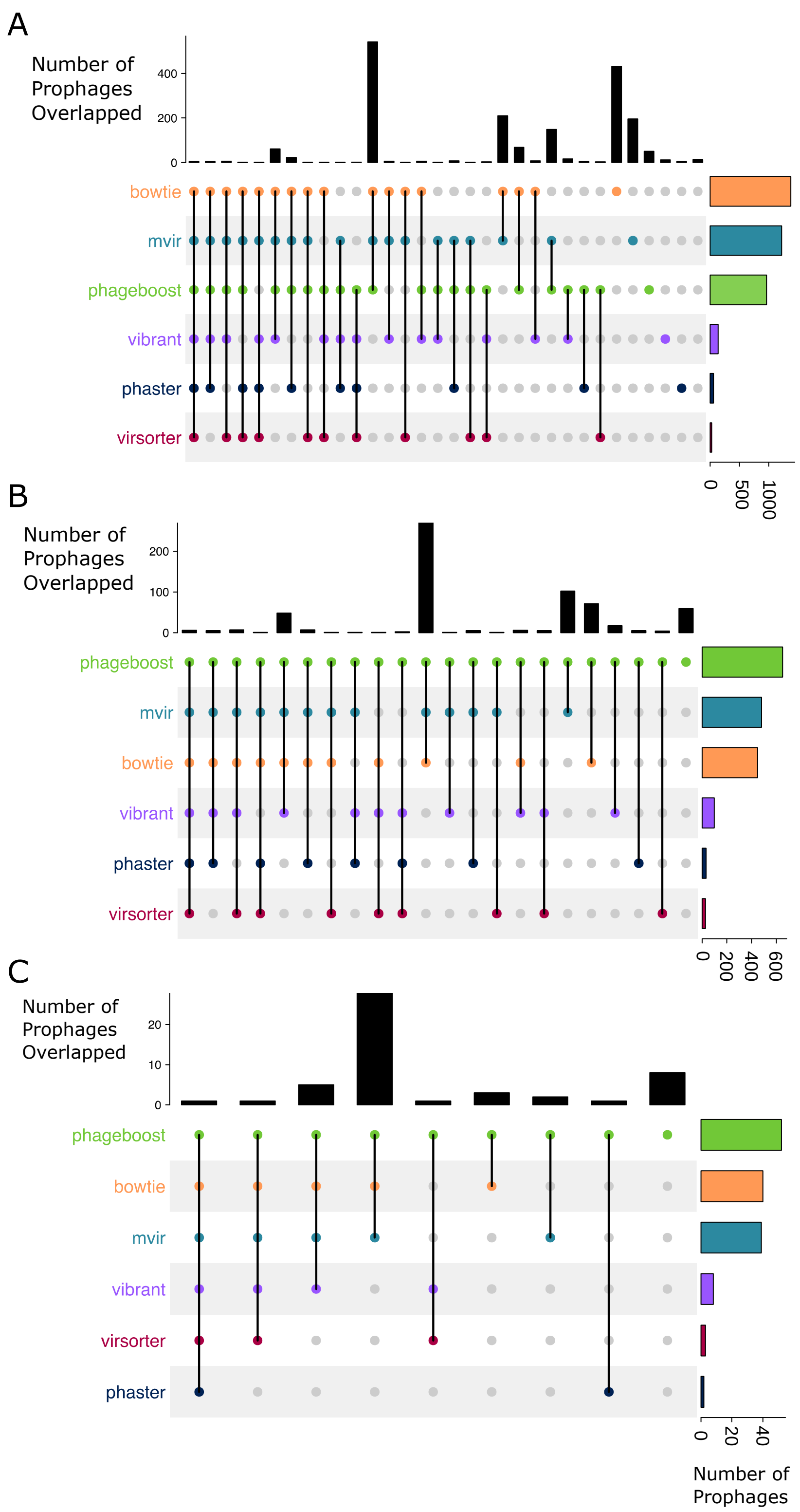

A
Number of Prophages Overlapped
0
200
400
bowtie
mvir
phageboost
vibrant
phaster
virsorter
0
500
1000
B
Number of Prophages Overlapped
0
100
200
phageboost
mvir
bowtie
vibrant
phaster
virsorter
0
200
400
600
C
Number of Prophages Overlapped
0
10
20
phageboost
bowtie
mvir
vibrant
virsorter
phaster
0
20
40
Number of Prophages

**Supplementary Figure 2 Upset Plot Showing Overlap Between Prophage Predictors:**

Summary of prophage predictions by Bowtie, mvir, phageboost, vibrant, phaster, virsorter of (A) All the 2, 719 merged prophage regions (B) All the 651 prophages that meet our phage criteria (C) and the 52 active prophages

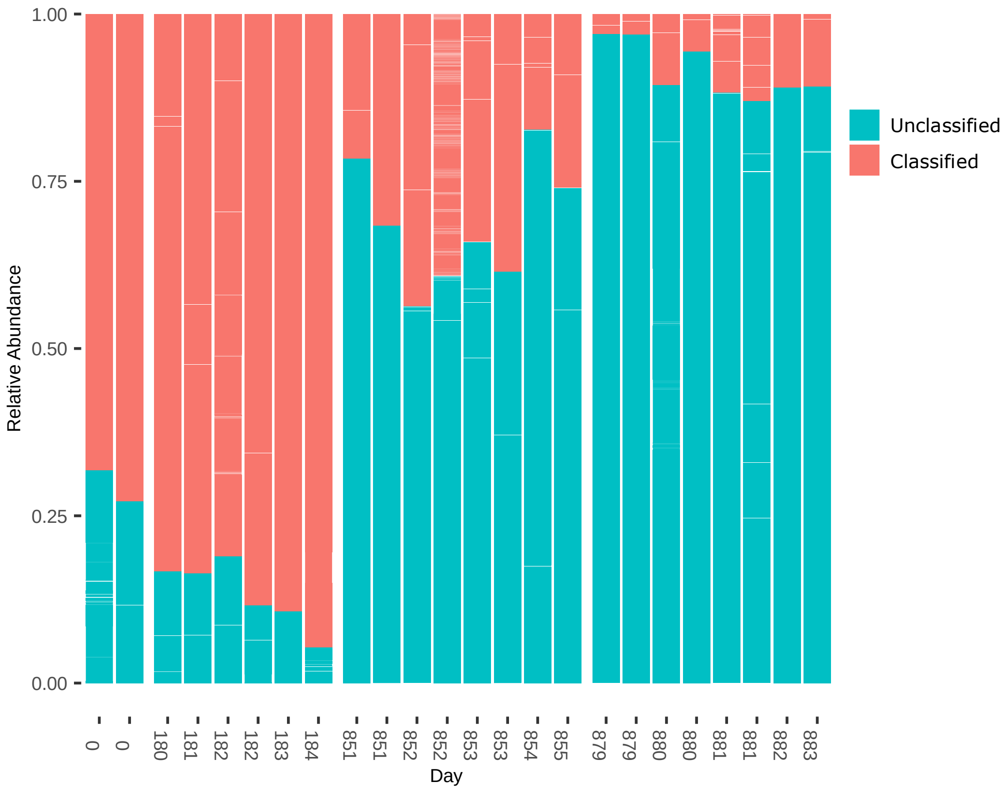


**Supplementary Figure 3 Percentage of Phages with Taxonomic Classification at the Family-Level per Sample-Sequence Run:**

Percentage of phages with taxonomic classification at the family-level per sample-sequence run. Individual contigs are separated by grey-lines showing the breakdown of individual contigs.
